# Trustworthy ML/AI for Aging Clocks: Preventing Systematic Prediction Bias in Biological Age Estimation

**DOI:** 10.64898/2026.05.27.728155

**Authors:** Hwiyoung Lee, Zhenyao Ye, Yun Yang, Yezhi Pan, Bradley A. Maron, Ze Wang, Peter Kochunov, Paul M. Thompson, L. Elliot Hong, Tianzhou Ma, Chixiang Chen, Shuo Chen

**Author notes:** Contributing authors. These authors contributed equally to this work (Hwiyoung Lee, Zhenyao Ye).

## Abstract

Machine learning (ML)- and artificial intelligence (AI)-based aging clocks are increasingly used to quantify physiological and molecular aging from omics and medical imaging data as distinct from chronological age. Here, we characterize a fundamental but underappreciated statistical limitation of commonly used ML/AI regression models for continuous outcomes: systematic prediction bias and its propagation to downstream association estimates. This issue becomes more challenging when the true outcome, biological age, is latent and therefore unobserved during ML/AI model training. We demonstrate that systematic prediction bias can distort and, in some cases, even reverse downstream association analyses that use aging clocks as ML/AI-predicted outcomes to assess their associations with exposures or clinical factors. For example, it can produce spurious associations suggesting that older predicted brain age is linked to better cognitive performance, or that older epigenetic age is associated with better kidney function. To address this problem, we introduce a principled and broadly applicable ML/AI regression framework based on constrained optimization, yielding better calibrated aging-clock estimates and valid downstream inference.

---

An aging clock (Figure 1) is a predictive tool, often derived from omics or imaging data, that estimates biological age relative to chronological age [1]. Biological age reflects the physiological and molecular characteristics of aging and captures how “old” an individual is at the biological level [2, 3]. This metric can deviate from chronological age due to various genetic, environmental, lifestyle, and disease related factors. Accelerated aging, in which biological age exceeds chronological age, is often associated with an increased risk of age-related diseases or adverse health-related exposures[4–8].

**Fig. 1:**
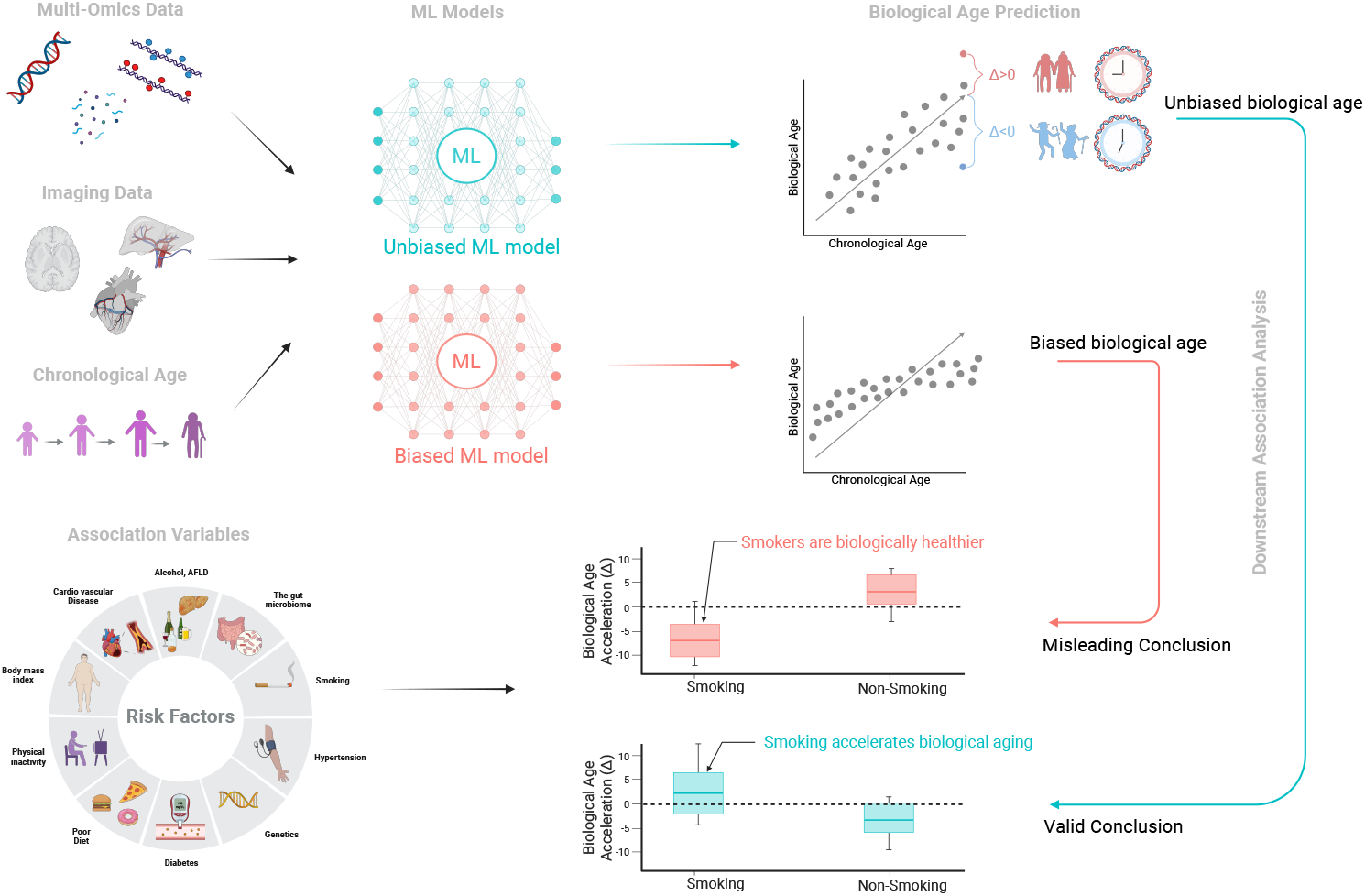
Illustration of biological age research and the influence of bias. Omics and imaging data collected from biological assays can be used to objectively assess biological age, which reflects an individual’s underlying aging status. Because biological age cannot be directly observed, ML/AI methods are commonly used to estimate it from omics and imaging features. An individual’s estimated biological age may differ from chronological age because of genetic, environmental, lifestyle, and disease-related factors. Using the estimated biological age, downstream analyses typically aim to identify exposures or clinical conditions associated with biological age resilience or acceleration and to assess whether these relationships may be causal. Here, resilience and biological age acceleration refer, respectively, to an estimated biological age that is younger (Δ *<* 0) or older (Δ *>* 0) than chronological age. However, the systematic prediction bias of commonly used ML/AI regression models can lead to systematically biased biological age estimation and thus misleading biomedical conclusions (as illustrated by the biased path: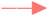). These biases can be prevented by developing unbiased ML regression models under an advanced computational framework (as illustrated by 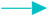)

Recent advances in aging clocks, including epigenetic, transcriptomic, proteomic, and neuroimaging-based approaches, have advanced aging research by helping to uncover mechanisms of aging, identify biomarkers of abnormal aging processes, and assess the effectiveness of preventive and therapeutic interventions for age-related diseases [9–15].

Instead of directly measuring an underlying biological age, aging clocks are computed indices derived from high-throughput multi-omics and imaging data using techniques such as machine learning (ML) and artificial intelligence (AI) regression models [16, 17]. These models use chronological age or age-related phenotypes as the outcome variable and omics or imaging features as predictors to approximate the *unknown* biological age [3, 10]. However, as noted by [18, 19], standard ML regression methods (e.g., random forests, gradient boosting, neural networks, and regularized regressions) often exhibit systematic prediction bias by shrinking predictions toward the mean. As a result, predicted biological ages tend to be underestimated at the upper end of the age range and overestimated at the lower end, in both training and testing sets (Figure 2). This bias is ‘systematic’, because prediction error, defined as the difference between predicted biological age and chronological age, varies systematically with chronological age rather than arising randomly. Although such bias is often treated using post hoc calibration or correction strategies, these approaches can make predicted biological age overly dependent on chronological age, thus limiting its ability to capture biological variation beyond age itself and making it less suitable as a measure of biological aging [20].

**Fig. 2:**
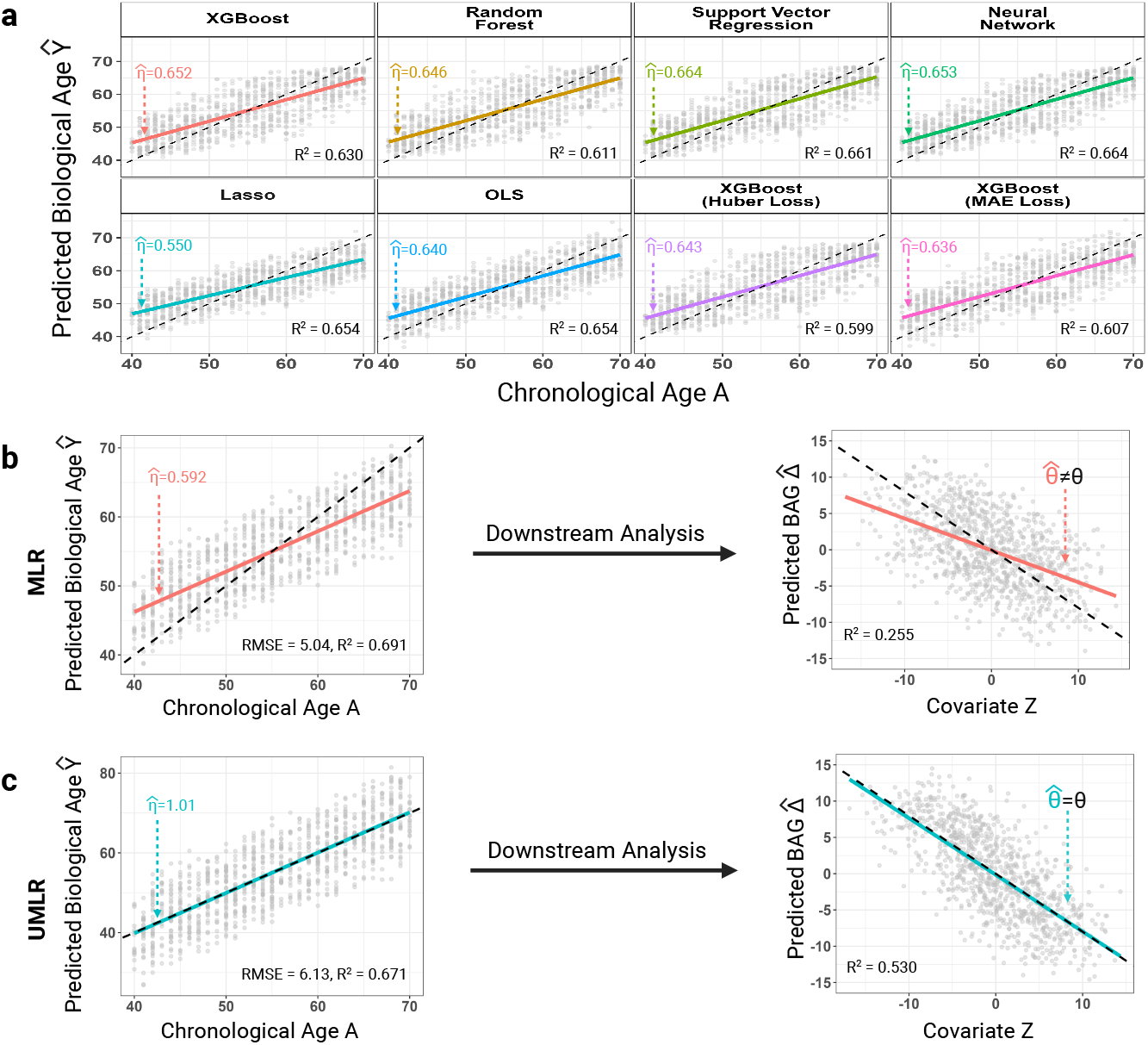
Illustrative synthetic experiment: systematic prediction bias in aging clock estimation from machine-learning (ML) and AI regression models, and its impact on downstream analyses. systematic prediction bias is present for MLR-derived biological age across standard ML/AI regression models (e.g., random forests, XGBoost, neural networks, LASSO, and others). Scatter plots display the predicted outcome *Ŷ* versus the true outcome *Y*, where the dashed lines represent the no-bias reference and the solid lines represent the fitted regression slope,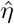. Systematic prediction bias is present when 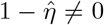; that is, when the solid line deviates from the dashed line. This bias is ubiquitous across the ML regression (MLR) models and objective functions considered here. (**b**) systematic prediction bias in general MLR models and MLR-derived biological age models affects downstream association analyses, introducing biased estimates of the association between covariates and the biological age gap and potentially leading to mistaken conclusions. (**c**) systematic prediction bias is approximately zero under the general unbiased MLR (UMLR) model and in UMLR-derived aging clock applications, leading to unbiased estimates of the biological age gap and its associations with covariates.

More importantly, the focus of many biological age studies is to identify associations between health-related factors and biological age, which is often referred to as *inference with predicted outcomes* in the recent literature [21–23]. The systematic prediction bias in ML/AI-predicted biological age can lead to biased downstream association estimates and, in causal settings, biased causal effect estimates. Such distortions may, in turn, misinform public health and clinical decisions, posing a significant challenge for studies that rely on aging clocks. These serious consequences have received far less attention in biological-age applications, partly because true biological age is latent, leaving no gold-standard measurements for training or validating predictions.

In this work, we address this issue by developing a statistical framework for quantifying and correcting systematic prediction bias in aging clocks. We first introduce a metric that quantifies the systematic prediction bias in aging clocks induced by ML/AI regression models. We further establish a quantitative relationship characterizing how this bias propagates into downstream estimates of associations between accelerated biological aging and health-related exposures or disease outcomes. Together, these findings underscore the importance of assessing and preventing systematic bias in aging clocks to ensure valid biomedical inference. To address this problem, we develop an unbiased ML/AI-based aging clock framework that directly prevents systematic prediction bias during model construction.

## Results

We begin by analytically characterizing systematic prediction bias as a prevalent phenomenon in machine-learning regression (MLR). We then distinguish biological-age applications from standard MLR settings: in biological-age applications, true biological age is latent, and chronological age or related variables serve as surrogate outcomes for model training, whereas in standard MLR settings, the outcome of interest is directly observed.

### Systematic prediction bias in general MLR

Let **X**_*i*_ *∈* ℝ^*p*^ denote the vector of *p* high-throughput predictors for subject *i*, and *Y*_*i*_ denote the corresponding outcome, for *i* = 1, *· · ·*, *n*. The training data are given by **X** = (**X**_1_, *· · ·*, **X**_*n*_)^*⊤*^ *∈* ℝ^*n×p*^ and **Y** = (*Y*_1_, *· · ·*, *Y*_*n*_)^*⊤*^ *∈* ℝ^*n*^. Without loss of generality, we center the outcome so that 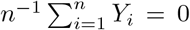. Let *f* denote an MLR predictive function, and define the predicted value for subject *i* as *Ŷ*_*i*_ = *f* (**X**_*i*_).

### Prediction bias

Prediction bias can be assessed by comparing the response variable **Y** with its predicted value Ŷ (see Fig S1. in *SI*). The predicted outcome is unbiased if E(*Ŷ*_*i*_ | *Y*_*i*_ = *y*^*∗*^) = *y*^*∗*^ for all *y*^*∗*^ in the support of *Y*, yielding a 45-degree line for predicted outcomes Ŷversus outcomes *Y*.

Let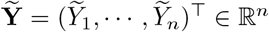 denote biased predictions in the sense that 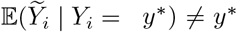. Throughout, Ŷ denotes a generic prediction, while 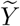 denotes a prediction that is specifically biased. The prediction bias is ‘*systematic*’, if the bias depends on the true outcome through a deterministic function *h*(·), that is, 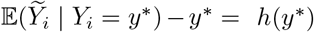.

As pointed out by [19] and illustrated in *SI*, systematic prediction bias arises from optimizing commonly used MLR objective functions (e.g., MSE or Huber loss), which are shared across many machine learning regression models (e.g., shrinkage regression, random forest, and neural networks).

Based on extensive numerical evidence and Theorem 2 in *SI*, a common form of systematic prediction bias is linear shrinkage toward the mean, 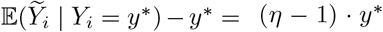, where *η ∈* [0, 1]. We quantify the degree of systematic prediction bias by 1*− η* (or 100(1 *−η*) %), with 1 *−η* = 0 indicating no bias. Under the linear bias pattern, *η* reflects the extent of systematic prediction bias through its attenuation of the Ŷ and *Y* relationship, with bias most pronounced at the extremes, where larger outcomes tend to be systematically underestimated and smaller outcomes tend to be systematically overestimated. In practice, *η* can be estimated as the slope coefficient from regressing Ŷ on **Y**.

#### Why does MLR yield biased predictions ?

MLR procedures may favor systematically biased predictions because they can attain a lower value of the training objective (see *Proposition* 1 in [19]). For example, under an MSE objective, biased predictions 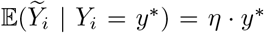 can achieve a smaller training loss than unbiased predictions E(*Ŷ*_*i*_ | *Y*_*i*_ = *y*^*∗*^) = *y*^*∗*^, i.e.,

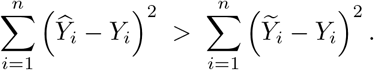

This can be viewed as a special case of the bias-variance trade-off: introducing linear systematic prediction bias can reduce variance and thereby lower the MSE. In settings where the primary goal is to minimize overall prediction error, such bias may be acceptable. However, in aging-clock applications, systematic prediction bias can distort biological-age estimates and lead to misleading downstream inferences relevant to healthy aging.

#### Bias in MLR-derived biological age

In aging-clock applications, the target quantity, underlying biological age *Y*, is *unobserved*, unlike in standard MLR settings where the outcome is directly observed. A common strategy is to train ML/AI models using chronological age *A* as a surrogate outcome and to treat the resulting prediction Â as a predicted biological age, denoted *Ŷ*. The rationale is that, at the population level, biological age is intended to be calibrated to chronological age within each chronological-age group, i.e., E(*Y*_*i*_ | *A*_*i*_ = *A*^*∗*^) = *A*^*∗*^. Accordingly, a well calibrated aging clock should satisfy E(Ŷ_*i*_|*A*_*i*_ = *A*^*∗*^) = *A*^*∗*^. In practice, however, systematic prediction bias is frequently observed in aging-clock applications (Figure 2 **a**), violating this requirement, with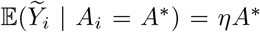, where *A* is assumed to be centered 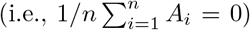 without loss of generality.

This bias is inherited directly from the systematic prediction bias in general MLR because Â is a biased prediction of *A*. Although the true biological age is unobserved, the degree of systematic prediction bias, *η*, can still be estimated by regressing **Ŷ** on **A**. We show that the resulting estimator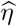 converges to the value that would be obtained if the latent biological age were observed (see Theorem 1 in *SI*). Therefore, we adopt the same approach used earlier to quantify the degree of bias in MLR-derived biological age as *η*.

#### Measuring systematic prediction bias for biological age applications by 1 − η

Despite the outcome being unobserved in MLR-derived biological-age applications, systematic prediction bias can still arise because the training objective (e.g., MSE) favors regression-to-the-mean shrinkage. Given the requirement that E(*Ŷ*_*i*_ | *A*_*i*_ = *A*^*∗*^) = *A*^*∗*^, unbiased prediction implies that the regression slope of *Ŷ* on *A* equals1. In the following Theorem 1, we show that the slope parameter 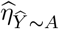 estimated from regressing *Ŷ* on *A* converges to 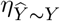, the corresponding slope that would be obtained if the true biological age *Y* were observed. This result shows that the bias measured with respect to chronological age reflects the underlying bias with respect to latent biological age. Thus, this result motivates our UMLR strategy: although *Y* is unobserved, enforcing 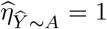 during model training ensures 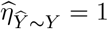, thereby mitigating systematic prediction bias relative to the latent outcome.

Without loss of generality, we first assume that chronological age *A*, true biological age *Y*, and biased predicted biological age 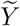 are all mean-centered, i.e.,

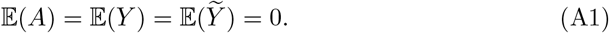

finite second moments;

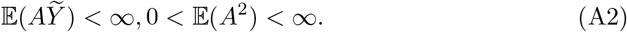

We suppose that the aging clock 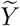 is a linearly, systematically biased with respect to the biological age *Y*, i.e.,

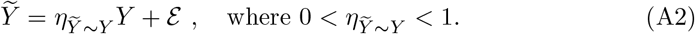

We also assume the standard mean zero residual condition, i.e.,

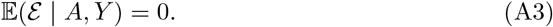

This assumption implies that, conditional on (*A, Y*), the residual ℰ has no systematic trend on average. We further consider that the chronological age is the baseline (i.e., normal) aging trajectory. Thus, at the population level, biological age follows chronological age, i.e., the average biological age among people of chronological age *A* = *a* equals *a*, i.e., for all *a*

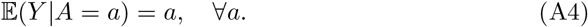

Let 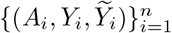 be independent identically distributed (i.i.d) observations, where 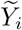 is the predicted biological age for subject *i* obtained from a pre-trained aging clock MLR model. In the following Theorem, we show taht the fitted regression slope 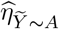 of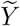 on *A* also converges in probability to 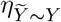.

##### Theorem 1

*Under assumptions (A1)–(A4), for biological age applications with surrogate outcome A, the estimated systematic prediction bias, given by the regression slope of* 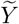 *on A, is a consistent estimator of the systematic prediction bias based on the observed ‘oracle’ biological age Y*; *namely*, 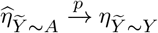*as sample size n → ∞*.

#### Impact on downstream biological age analyses

A primary goal in aging-clock studies is often to assess associations between biological age acceleration (gap), that is, predicted biological age subtracting the chronological age and risk factors/clinical conditions to inform interventions for healthy aging. Let Δ = *Y −A* denote the biological age gap (BAG). As shown below, systematic prediction bias in aging clocks can propagate into downstream association analyses, because biased biological-age predictions lead to biased estimates of BAG.

Let *θ* = cov(Δ, *Z* | **V**) denote the conditional covariance between the aging gap Δ and an exposure *Z* while controlling for confounders **V**. Let 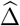 be an unbiased estimate of Δ, and let 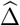 be a biased estimate from MLR [e.g., 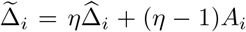, with 0 *< η <* 1), then

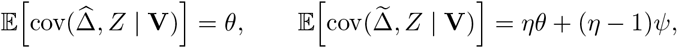

where E[cov(*A,Z* | **V**)] = *Ψ*. Thus, the bias of estimated association, 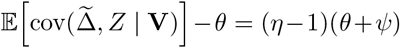, is generally nonzero. This bias can substantially affect biomedical conclusions because it can distort, or even reverse, the sign of the relationship between biological age and exposures or health conditions (see Figure 2 **b** and the case study results).

### A principled framework for preventing systematic prediction bias

Our goal is to prevent systematic prediction bias during the model training stage, ensuring *η* = 1. To achieve this, we implement the following constraints on the MLR objective function:

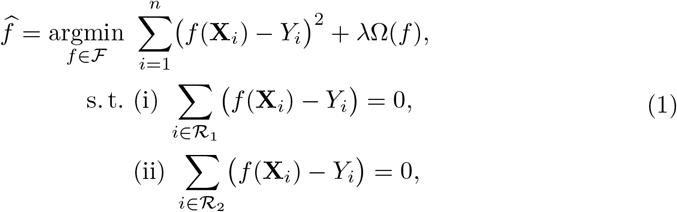

where Ω is a regularizer and *λ* is a tuning parameter, which controls model complexity to mitigate overfitting and improve generalization. *ℛ*_1_ and *ℛ*_2_ denote two disjoint subsets of participants whose outcome values lie in complementary, continuous supports (i.e., two disjoint subsets of the training data defined by a cutoff *Y*_*c*_), *ℛ*_1_ = *{i* : *Y*_*i*_ *≤ Y*_*c*_*}* and *ℛ* _2_ = *{i* : *Y*_*i*_ *> Y*_*c*_*}*, for example, 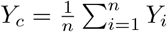.

These constraints calibrate the prediction function in two disjoint outcome regions, thereby eliminating systematic prediction bias and preventing slope attenuation, as supported theoretically by Theorem 2 in *SI* showing the systematic prediction bias having a linear pattern and Theorem 1 in [19] showing that the constraints correcting the linear bias.

We refer to this strategy as unbiased MLR (UMLR). As demonstrated in Figure 2 and the simulations in *SI*, UMLR removes systematic bias in both general MLR settings and MLR-derived aging clocks. Importantly, association estimates between biological age gaps and risk factors obtained using UMLR are unbiased (Figure 2 c-right), which is critical for biological age and healthy aging research.

#### A simple explanation of UMLR method

The commonly observed linear pattern of systematic prediction bias for conventional MLR in the Ŷ –*Y* relationship can be characterized by a line passing through a single point (for example, 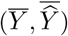). However, this alone does not guarantee that the slope of Ŷ on *Y* equals 1 (that is, *η* = 1, or equivalently 1 *− η* = 0, indicating no bias).

Geometrically, two distinct points determine a unique line (see Figure 3). Motivated by this principle, UMLR uses two mean matching constraints to define two anchor points in two disjoint outcome regions, ℛ_1_, and ℛ_2_). Under the assumed linear bias pattern, the constraints enforce the unbiased 45-degree line, implying *η* = 1. When the distribution of *Y* is known, the cutoff defining the constraint subsets can be chosen optimally to improve the effectiveness of the constraints, for example by achieving better subgroup separation and more balanced subgroup sizes. In practice, however, this information is often unavailable. We therefore use the sample mean of the outcome in the training set as the default cutoff for defining the two subsets. Under this principle, unbiased prediction holds when UMLR uses multiple constraint subsets that together span the full range of the outcome.

**Fig. 3:**
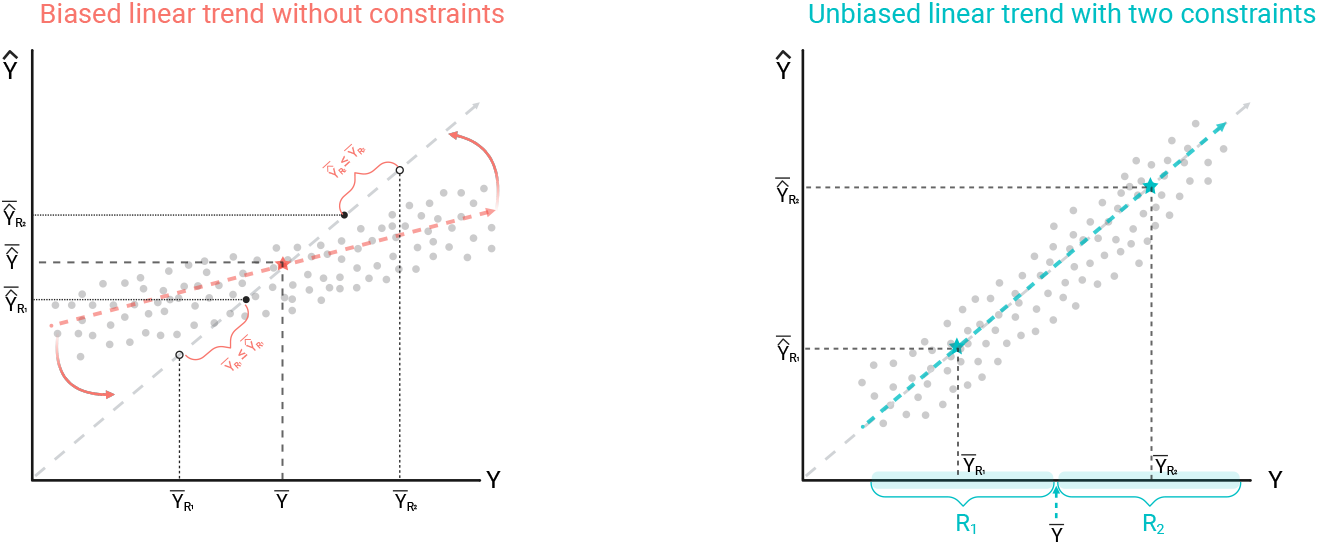
This figure illustrates the intuition behind UMLR for achieving unbiased prediction. Conventional MLRs (left) enforce that the fitted line passes through 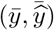, but they do not guarantee a slope of 1. In contrast, UMLR (right) imposes constraints that force the fitted line to pass through two distinct points on the unbiased prediction line. Geometrically, these two anchoring points uniquely determines the line, thereby ensuring that the fitted line coincides with the unbiased prediction line and has slope 1.

As shown in Figure 2 **c**, the unbiased *Ŷ* produced by UMLR can have a larger MSE and, consequently, a lower *R*^2^ between predicted biological age and chronological age than conventional MLR models (Figure 2 **b**). However, because biological age is unobserved, *R*^2^ can only be interpreted as concordance with the surrogate outcome (i.e., chronological age) rather than as an accuracy measure for the true biological age. In the extreme case, an *R*^2^ close to 1 would imply that biological age is nearly identical to chronological age for all individuals, providing little information beyond chronological age. Accordingly, a moderately lower *R*^2^ from UMLR may indicate that the clock is less constrained to mirror chronological age and can better capture biologically meaningful deviations from chronological age. Indeed, in simulation studies (*SI*), UMLR-derived estimates are more closely aligned with the true biological age than those from conventional MLR. More importantly, as shown in Figure 2, compared with the traditional MLR-derived BAG, the UMLR-derived BAG yields unbiased association estimates and achieves a higher *R*^2^ with the exposure *Z*, providing a more biologically informative aging measure and enabling more accurate downstream inference.

### Biological age case studies

In biological age research, brain age and epigenetic aging clocks are two most widely used applications that leverage neuroimaging and DNA methylation (DNAm) data to objectively measure underlying biological age. Here, we calculate brain age and epigenetic clock measures in two case studies, and evaluate systematic prediction bias in biological age estimation and its impact on downstream analyses.

#### Case Study 1

We constructed a brain-age clock by training models on neuroimaging data from the UK Biobank (UKB) and evaluating the trained models on the Human Connectome Project in Aging (HCP). The UKB is a large prospective study that recruited approximately 100,000 participants who underwent magnetic resonance imaging, generating multimodal brain imaging data that cover structural, functional, and diffusion modalities [24]. The HCP cohort recruited over 1,200 cross-sectional participants who also underwent multimodal brain imaging scans [25]. Previous neuroimaging studies have established that brain white matter (WM) integrity is highly reliable measure and associated with cognitive functions, and thus particularly suited for studying brain aging [6, 7, 26–33]. Therefore, we focused on brain WM structural integrity, measured by fractional anisotropy (FA), which was available in both cohorts. The analytic sample included 36,812 participants from UKB (mean (SD) age = 63.41 (7.52) years; age range = [45.00, 82.00], 54.95% female) and 391 participants from HCP (mean (SD) age = 60.47 (9.64) years; age range = [45.17, 79.67], 57.03% females).

We compared standard MLR methods with UMLR, and Figure 4 summarizes the estimated bias parameter *η*. As shown in Figure 4, the systematic prediction biases 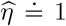 for brain age derived by UMLR suggest unbiased estimates in both training testing data set. In contrast, the systematic prediction bias *η* persists across all classic MLR in training and testing, overestimating biological age for younger participants and underestimating for the older.

**Fig. 4:**
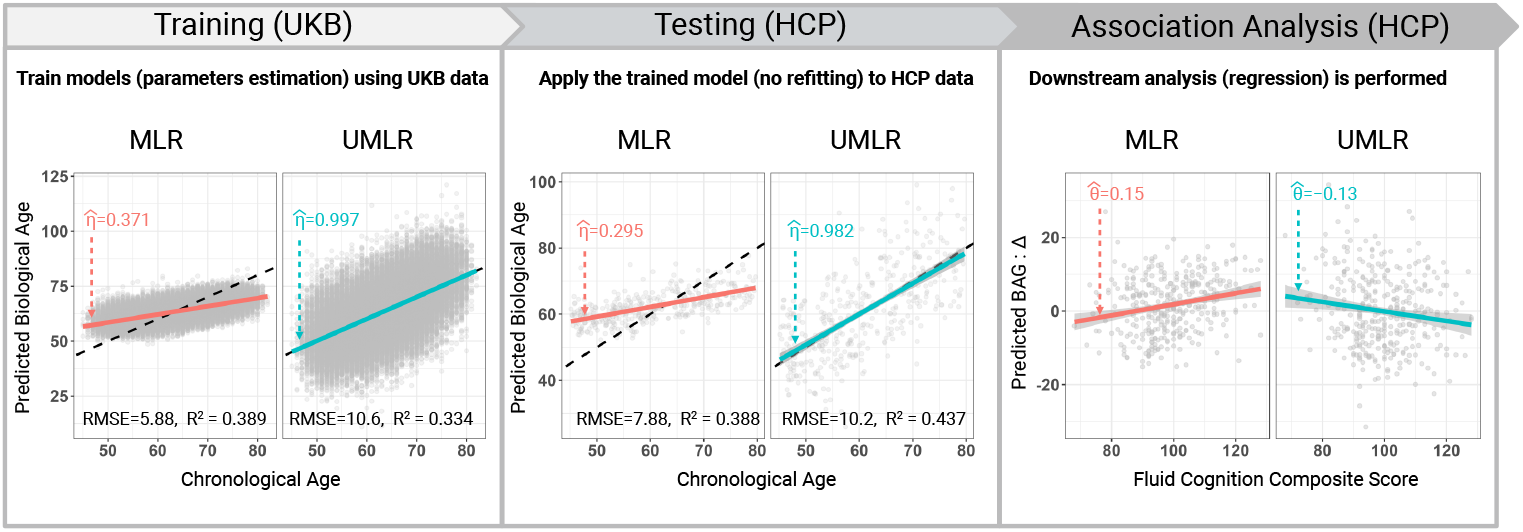
Case study 1. Brain-age analysis using neuroimaging data from the UK Biobank (UKB) as the training set and the Human Connectome Project (HCP) as an independent test dataset. Both conventional machine-learning regression (MLR) models and the proposed unbiased machine-learning regression (UMLR) were used to derive brain age. We characterize systematic prediction bias using scatter plots of predicted biological age versus chronological age, where the dashed lines represent the no-bias reference and the solid lines represent the fitted regression slope,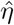. Systematic prediction bias is present when 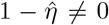; that is, when the solid line deviates from the dashed line. In the training set, conventional MLR (solid line in 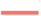) exhibits systematic prediction bias in exchange for higher *R*^2^ and lower MSE, whereas UMLR prevents this bias. In the test set, the same systematic prediction bias persists for MLR, whereas UMLR (solid line in 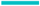) remains unbiased while achieving a higher *R*^2^. This bias in brain-age estimation can have a substantial impact on the down-stream analysis. Specifically, the brain-age gaps derived from UMLR and MLR yield *opposite* associations with the fluid cognition composite score: fluid cognition is negatively associated with the UMLR-derived brain-age gap (i.e., higher fluid cognition corresponds to a more resilient, younger brain age), but positively associated with the commonly used MLR-derived brain-age gap (i.e., higher fluid cognition corresponds to a more accelerated, older brain age). Overall, using conventional MLR and UMLR in biological-age applications can lead to contradictory scientific conclusions and, consequently, may imply different health-related decisions.

The association between calculated biological age and clinical or exposure variables can also be influenced by systematic prediction bias. However, unlike the simulation setting, the underlying true association is unknown in the real data. We therefore examined the association between estimated brain age and cognitive performance in the HCP test dataset. We used the Fluid Cognition Composite Score from the NIH Toolbox Cognition Battery. This battery is designed to assess multiple cognitive domains, including executive function, attention, episodic memory, language, processing speed, and working memory [34]. Individuals with lower composite scores have poorer cognitive functioning, and thus worse brain health.

As demonstrated in Figure 4, the UMLR-derived biological age gap was negatively associated with Fluid Cognition Composite Score (*t*_*df*=355_ = 4.38, *p* = 1.58 *×*10^*−*5^) after adjusting for covariates(e.g., biological sex), consistent with the expectation that accelerated brain aging is linked to worse cognition. In contrast, the traditional MLR-derived biological age gap was positively associated with cognitive performance (*t*_*df*=355_ = −2.62, *p* = 0.066), suggesting the opposite conclusion. These discordant results underscore that whether systematic prediction bias is addressed in ML/AI regression models can fundamentally alter, and even reverse, inferences in downstream analyses.

#### Case Study 2

Epigenetic clocks measure biological age based on DNA methylation patterns and have gained popularity in aging research due to their reliability and accuracy [15]. In the second case study, we built an epigenetic clock by training models on processed DNA methylation data from the Framingham Heart Study (FHS) and investigated the association between the epigenetic clock gap and serum creatinine concentration, an index for kidney function. Specifically, we used DNAm data from 1,476 participants collected at the University of Minnesota (UMN) as the training set, and data from 226 participants collected at Johns Hopkins University (JHU) as the testing set. A total of 2,990 CpG sites were used as predictors, including CpG sites from established epigenetic clocks [11, 35, 36] and additional sites selected through a sure independence screening procedure [37]. For demonstration purposes, we used LASSO to estimate biological age, although similar performance was observed with other regression methods. We then evaluated the association between the biological age gap and serum creatinine concentration in the testing data.

The sample demographics are summarized as follows. The training set included 1,476 UMN subjects (mean (SD) age = 65.12 (8.71) years; age range = [40, 92]; 54.40% female), and the testing set included 226 JHU subjects (mean (SD) age = 70.32 (8.50) years; age range = [49, 91]; 18.58% female). Despite these demographic differences, biological age methods show consistent performance in the training and testing sets (see Fig 4).

Figure 5 illustrates biological age estimation performance in the training and testing sets for both MLR and UMLR. No systematic prediction bias was observed for UMLR, with 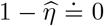 in both the training and testing sets. In contrast, systematic prediction bias was present for MLR in both sets, with 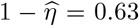 in the training set and 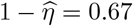 in the testing set. MLR achieved better performance in terms of *R*^2^ and RMSE (training set: *R*^2^ = 0.49, RMSE = 6.41; testing set: *R*^2^ = 0.42, RMSE = 7.48), which reflects the variance-bias trade-off in machine learning and statistics. In the downstream analysis, we tested the association between epigenetic clock-derived biological age gap and serum creatinine concentration (mg/dL), a marker of kidney function reflecting glomerular filtration and creatinine clearance. The serum creatinine concentration at a higher level suggests a worse health condition, which may be correlated with an accelerated epigenetic aging clock (i.e., positive biological age gap). However, as demonstrated in Figure 5, the biological age gap derived by conventional MLR yields a negative adjusted association (i.e., younger epigenetic clock is linked with higher creatinine concentration) with *t*_*df*=220_ = −3.68, *p* = 2.94 *×*10^*−*4^. In contrast, a positive association between biological age gap and creatinine concentration was estimated using UMLR (*t*_*df*=220_ = 0.79, *p* = 0.43) was obtained by biological age gap using UMLR. These results again demonstrate the influence of systematic prediction bias in downstream association analysis.

**Fig. 5:**
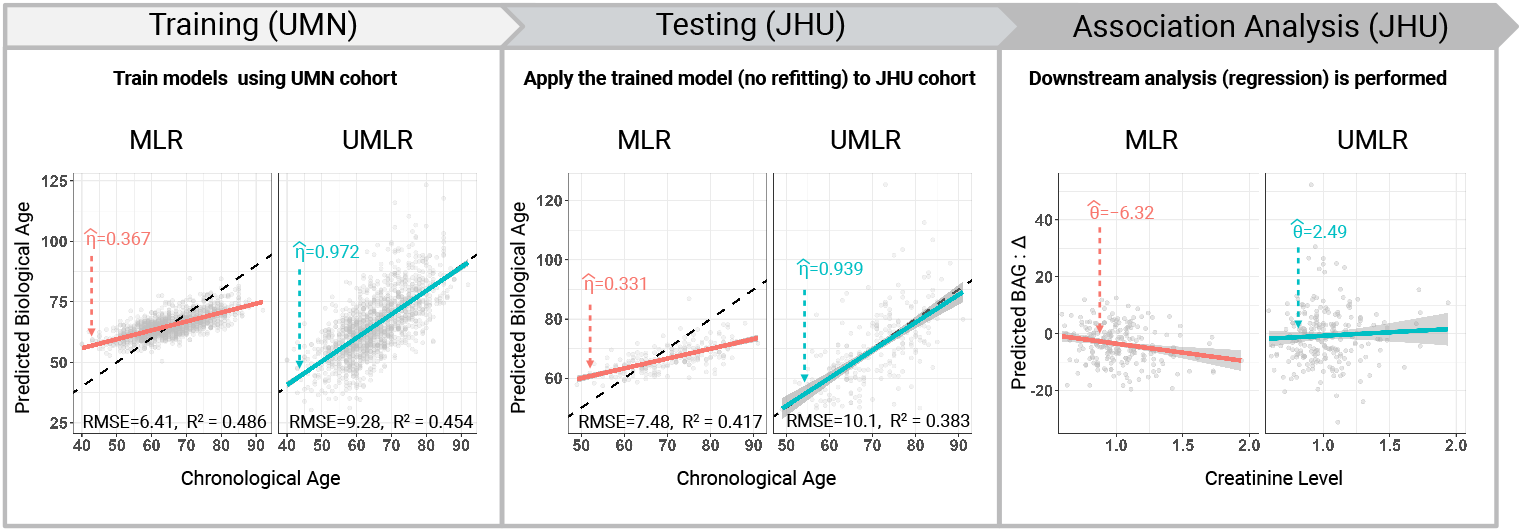
Case study 2: Epigenetic aging clocks were constructed from DNA methylation (DNAm) data in the Framingham Heart Study cohort. DNAm data from the University of Minnesota (UMN) site were used for training, whereas data from the Johns Hopkins University (JHU) site were used for testing. Systematic prediction bias was evident for epigenetic clocks estimated using conventional MLR in both the training and testing sets (solid line in 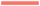). By contrast, the systematic prediction bias was approximately zero under the UMLR approach (solid line in 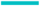). In downstream analyses of the testing set, the association between the epigenetic biological age gap and serum creatinine concentration, an indicator of kidney function for which higher concentrations indicate worse kidney function, reversed direction between MLR and UMLR, leading to *opposite* conclusions (i.e., worse kidney function was associated with a younger MLR-derived epigenetic age but an older UMLR-derived epigenetic age).

**Fig. 6:**
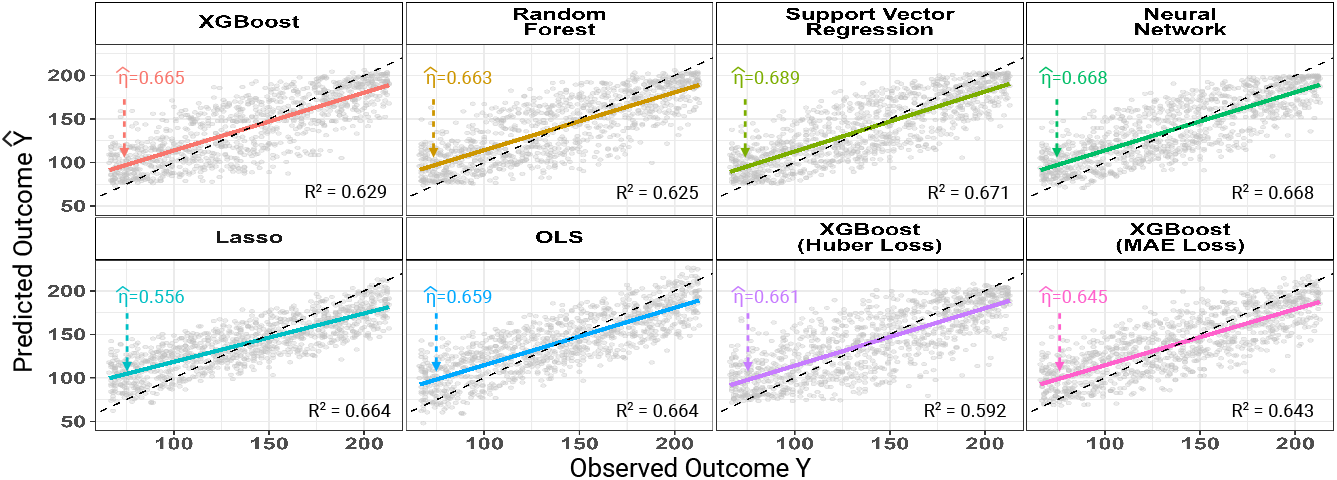
Illustrative synthetic experiment: systematic prediction bias in machine-learning (ML) and AI regression models. Systematic prediction bias is present across standard ML/AI regression models (e.g., random forests, XGBoost, neural networks, LASSO, and others). Scatter plots display the predicted outcome Ŷ versus the true outcome *Y*, where the dashed lines represent the no-bias reference and the solid lines represent the fitted regression slope, 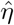. Systematic prediction bias is present when 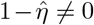; that is, when the solid line deviates from the dashed line. This bias is ubiquitous across the ML regression (MLR) models and objective functions considered here.

## Discussion

Systematic prediction bias, characterized by shrinkage of fitted values toward the outcome mean, is widely observed in machine-learning regression analyses, yet its implications are often overlooked when models are evaluated primarily by overall prediction error. Under commonly used training objective functions such as MSE, such bias may even be favored when shrinkage reduces variance sufficiently to improve aggregate predictive performance, consistent with the bias–variance trade-off. We therefore formally define systematic prediction bias and introduce measures that quantify it directly, rather than leaving it to be inferred from aggregate error metrics that are insensitive to its presence. In biological-age research, the true biological age is unobserved, and aging clocks are typically constructed by training MLR models that regress chronological age on omics or imaging features. Despite this key difference from standard MLR settings with observed outcomes, we extend our theoretical results to this setting and show that the proposed measures remain well defined and that the systematic prediction bias persists: predicted biological age deviates from chronological age in a systematic, frequently linear, pattern, even though the population mean biological age is expected to align with chronological age within each chronological-age stratum. Older individuals may thus appear to have artificially “resilient” biological age (i.e., substantially lower predicted biological age than chronological age) simply due to the linear shrinkage instead of underlying biological conditions. Such distortions further propagate to downstream analyses, which is of particular consequence because aging clocks have recently been used with increasing frequency as outcomes, primary endpoints, and biomarkers [12, 38–41]. The goal of these studies is often to evaluate the relationships between aging clocks and health conditions, treatments, or exposures, a task that falls within the framework of inference with predicted outcomes. However, our theoretical results show that such inference, when based on conventional ML/AI-regression-derived aging clocks, can yield biased estimates and misleading conclusions.

Here, we provide unbiased ML/AI regression frameworks as a general and principled solution to prevent systematic prediction bias by imposing additional constraints on conventional objective functions used in MLR. These constraints act at the parameter-optimization stage and yield parameter estimates that ensure unbiased prediction, making the approach applicable to both general ML/AI regression models for continuous outcomes and biological-age prediction. In biological-age applications, UMLR can better characterize biological age because (i) it corrects bias, particularly in younger and older participants, and (ii) it more accurately captures deviations of biological age from chronological age. More importantly, UMLR yields unbiased down-stream association estimates between biological-age measures and covariates, which are crucial for informing medical and public health decisions.

### Software

A web application is provided at https://github.com/hwiyoungstat/UMLR to demonstrate the systematic prediction bias in traditional AI/ML regression models, and illustrative how this bias leads to biased aging clocks and downstream association analyses. We also provide examples of unbiased machine learning regression (UMLR) models and user-friendly `R` functions for implementing UMLR.

## UK Biobank

This research has been conducted using the UK Biobank Resource under application number 74376.

## Framingham Heart Study

From the Framingham Heart Study of the National Heart Lung and Blood Institute of the National Institutes of Health and Boston University School of Medicine. This project has been funded in whole or in part with Federal funds from the National Heart, Lung, and Blood Institute, National Institutes of Health, Department of Health and Human Services, under Contract No. 75N92019D00031.

## Supplementary Information

### Proof of Theorem 1

*Proof*

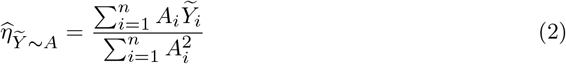

By the weak law of large numbers (WLLN), 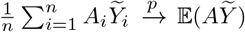 and 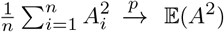

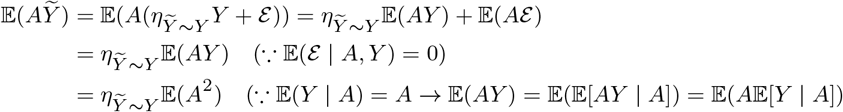

By the continuous mapping theorem, we obtain 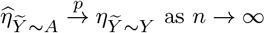.

### Linear Relation Between ML/AI Predicted Outcome and the True Outcome

#### Theorem 2

(Linear Relation Between ML/AI Predicted Outcome and the True Outcome) *Let Y ∈* ℝ *be the true outcome and let* 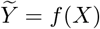 *be the prediction of any ML/AI regression model f ∈ ℱ trained to minimize the mean squared error* E [(*Y − f* (*X*))^2^] *over a restricted function class E. Assume further that* 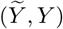*is jointly normal. Then*,

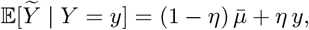

*where* 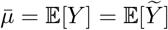 *and* 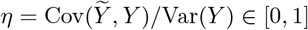.

*Proof*

#### 1. MSE minimization and the projection condition

Any ML/AI regression model trained under squared error loss solves:

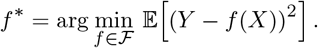

For any competitor *f*_*λ*_ = *f* ^*\**^ + *λh* with *h ∈ ℱ* and *λ ∈* ℝ, the optimality of *f* ^*\**^ requires:

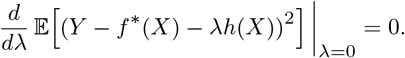

Expanding and differentiating:

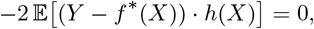

which gives the projection condition:

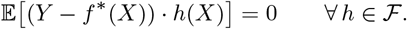

Two immediate consequences follow. Taking *h*(*X*) = 1 *∈ ℱ*:

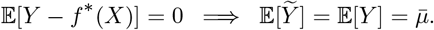

Taking *h*(*X*) = *f* ^*\**^(*X*) *∈ ℱ*:

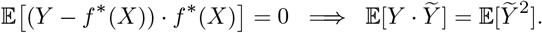

#### 2. Joint structure of 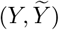

Write the prediction as 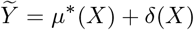, where *µ*^*\**^(*X*) = E [*Y* | *X*] is the oracle conditional mean and *δ*(*X*) = *f* (*X*) *− µ*^*\**^(*X*) is the misspecification error. Decompose the outcome as:

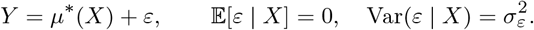

It follows that:

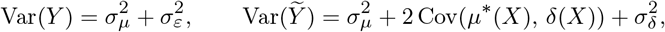

and:

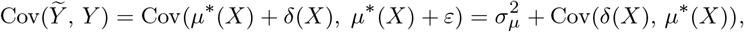

where 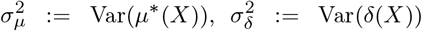, and the cross terms Cov(*µ*^*\**^(*X*), *ε*) = Cov(*δ*(*X*), *ε*) = 0 follow from *ε ⊥ X*.

#### 3. Deriving 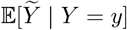

Applying the law of total expectation by conditioning on *X*:

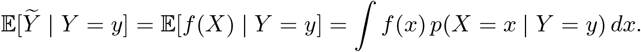

By Bayes’ theorem and *Y* = *µ*^*\**^(*X*) + *ε* with *ε ⊥ X*:

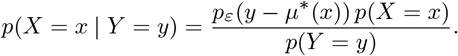

Substituting *f* (*x*) = *µ*^*\**^(*x*) + *δ*(*x*):

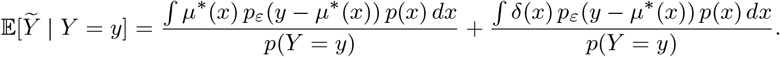

For the first term, make the change of variable *u* = *µ*^*\**^(*x*) and let *p*_*µ*_(*u*) denote the induced density of *µ*^*\**^(*X*):

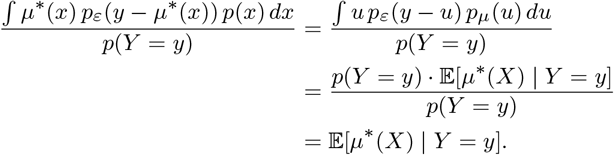

since *p*_*ε*_(*y − u*) *p*_*µ*_(*u*) = *p*(*Y* = *y, µ*^*\**^(*X*) = *u*). By the same argument the second term equals E [*δ*(*X*) | *Y* = *y*]. Therefore:

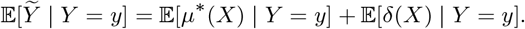

#### 4. Evaluating the conditional expectations under normality

Assume 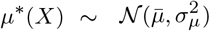 and 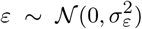 independently. Then the joint distribution of (*µ*^*\**^(*X*), *Y*) is bivariate normal:

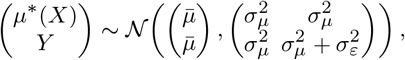

so the conditional expectation of *µ*^*\**^(*X*) given *Y* = *y* is linear in *y*:

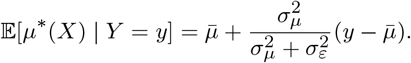

Similarly, assuming *δ*(*X*) is jointly normal with *Y* with Cov(*δ*(*X*), *Y*) = Cov(*δ*(*X*), *µ*^*\**^(*X*)) and E [*δ*(*X*)] = 0:

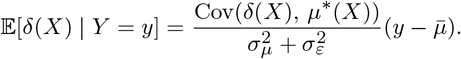

#### 5: Combining and identifying *η*

Adding the two conditional expectations:

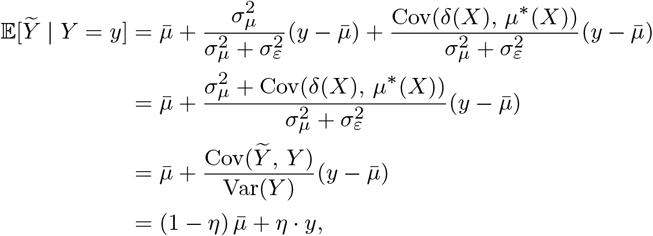

where:

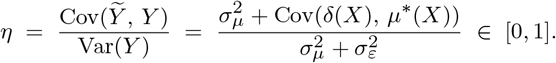

This establishes the linear relationship 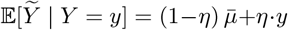 for any ML/AI regression model minimizing MSE over *ℱ*, under the normality assumption on (*µ*^*\**^(*X*), *δ*(*X*), *ε*). □

### Simulation Analysis

We conducted comprehensive simulation studies to evaluate systematic prediction bias in biological age estimates derived from MLR versus UMLR models and to assess how this bias propagates to downstream association analyses using biological age.

We simulated the synthetic data as follows. For each subject *i*, we first generated chronological age *A*_*i*_ from a uniform distribution with ages ranging from 40 to 100 years, i.e., *A*_*i*_ ~ Uniform(40, 100). We then generated an aging gap Δ_*i*_ ~ Uniform(*−*10, 10), and defined biological age as *Y*_*i*_ = *A*_*i*_ + Δ_*i*_. Given *Y*_*i*_, we generated a *p*-dimenaional predictor vector *X*_*i*_ *∈* ℝ^*p*^ as a function of biological age, so that *X*_*i*_ reflects biological age-related biomarker. Specifically, *X*_*i*_|*Y*_*i*_ *~ MV N* (`scale`(*Y*_*i*_), *σ***I**_*p*_), where scale 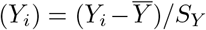. To evaluate how predicted biological age affects down-stream analyses, we additionally generated three external association variables that depend on biological aging gap (BAG) but are not used in the model fitting stage. Specifically, we generated two continuous exposure variables *Z*_1_, and *Z*_2_. *Z*_1_ is negatively correlated with BAG (i.e., corr(Δ, *Z*_1_) *≈ −*0.25), whereas *Z*_2_ is uncorrelated with BAG (i.e., corr(Δ, *Z*_2_) = 0).

We also generated a binary variable *D* for represents disease status as *D*_*i*_ = Bernoulli(*π*_*i*_), where *π*_*i*_ = logit^*−*1^(Δ_*i*_ *× β*). The disease risk is positively associated with Δ_*i*_: individuals with larger positive Δ_*i*_ (accelerated biological aging) have higher risk, whereas those with smaller Δ_*i*_ (biologically younger than expected given their chronological age) have lower risk. We repeated the simulation 100 times to evaluate the performance of biological age prediction and downstream analyses under UMLR and MLR. The results are summarized in Figure 7.

**Fig. 7:**
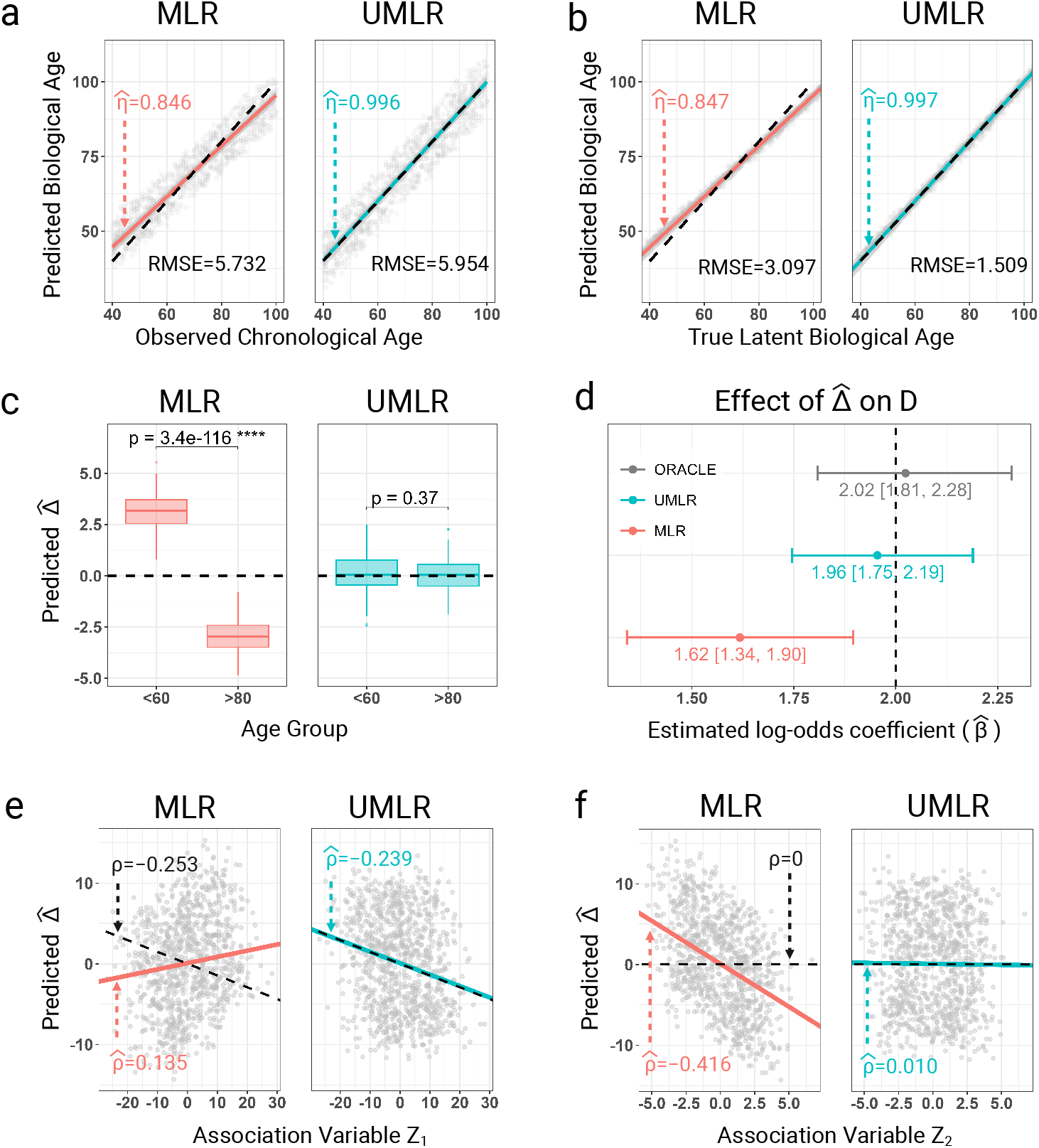
Simulation results: **a**, Systematic prediction bias for conventional MLR versus UMLR, comparing chronological age with predicted biological age in the testing set. Systematic prediction bias is present for MLR but not for UMLR, although MLR achieves a smaller RMSE. **b**, Systematic prediction bias for MLR versus UMLR, comparing true biological age with predicted biological age in the testing set. Systematic prediction bias is present for MLR but not for UMLR, while UMLR achieves a smaller RMSE. **c**, Comparison of the biological age gap between younger and older subcohorts; the dashed line denotes the ground truth. Systematic prediction bias in MLR artificially inflates apparent aging effects in older subcohorts (e.g., super-agers), a common source of misinterpretation in biological studies. **d**, Association analysis with disease as the outcome and biological age gap as the predictor. UMLR outperforms MLR, yielding more accurate estimates of the odds ratio OR. **e**, Association analysis with biological age gap as the outcome and exposure *Z*_1_ as the predictor (dashed line denotes the ground truth). The MLR-derived biological age produces an estimate with the incorrect sign, whereas UMLR provides accurate estimates. **f**, Association analysis with biological age gap as the outcome and exposure *Z*_2_, where no true association exists (dashed line denotes the ground truth). MLR yields a false-positive association, whereas UMLR provides unbiased estimates with parameter value 0.

#### Simulation results - Biological Age Prediction

We first evaluate the performance of biological age prediction. Specifically, we assess systematic prediction bias, quantified as 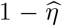, by regressing the predicted biological age on both observed chronological age and the true biological age. As shown in Figure 7 (**a,b**), MLR predictions exhibit clear systematic prediction bias 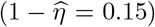, whereas UMLR shows essentially no systematic prediction bias, with 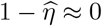.

Although UMLR yields a larger RMSE than MLR when evaluated against chronological age *A*, it achieves a smaller RMSE when evaluated against the true biological age *Y*. This result highlights that RMSE (or MAE) computed with respect to *A* alone may not be an appropriate metric for evaluating biological-age prediction, because it can differ substantially from the target metric, namely, RMSE with respect to *Y*. A more comprehensive evaluation of biological-age prediction should therefore also include systematic prediction bias.

#### Simulation results - Downstream Association Analyses

This systematic prediction bias in biological age can substantially influence downstream analyses, which are the primary focus of many biological aging studies. We consider three experimental settings to illustrate this impact.

In the first experiment, we compare the BAG between an elderly subgroup (Age *>* 80) and a younger subgroup (Age *<* 60). As shown in Figure 7 **c**, the MLR-derived BAG exhibited a negative mean in the elderly group and a positive mean in the younger group, yielding a false positive difference between the two groups, even though BAG was generated to be independent of age. In contrast, the UMLR derived BAG showed no evidence of a false positive group difference (*p* = 0.37).

In the second experiment, we treat the biological age gap as the predictor and disease status as the outcome and evaluate the odds ratio (OR). As shown in Figure 7 (**d**), the UMLR approach yields an unbiased estimate of the OR, whereas MLR produces a biased OR estimate.

In the third experimental setting, we consider the biological age gap as the outcome and exposures *Z*_1_ and *Z*_2_ as predictors. As shown in Figure 7 (**e,f**), the MLR approach yields biased estimates of the associations, while UMLR provides unbiased estimates. In Figure 7 (**e**), the direction of the association estimated by MLR is opposite to the ground truth (similar to the UKB case analysis), which can lead to misleading conclusions in practical studies. Finally, when no true association exists, the MLR approach can still falsely report statistically significant findings. Therefore, examining systematic prediction bias is essential prior to conducting downstream analyses.

### Materials for Case studies

#### Case Study 1: Brain age

In this study, we used data from two large cohorts, the UK Biobank and the Human Connectome Project in Aging. The UKB is a prospective cohort study with deep imaging phenotyping of approximately 100,000 participants aged 44 to 83 years across the United Kingdom [42]. The HCP provides rich neuroimaging data from over 1,200 healthy participants aged 36 to 100+ years [25].

We focused on WM brain age by using WM integrity measured by FA, derived from diffusion magnetic resonance imaging (dMRI) in both cohorts [6–8, 29, 43]. FA quantifies the directional dependence of water diffusion (range 0-1), with higher values reflecting greater microstructural organization and integrity of WM fiber tracts. We restricted the analysis to a common set of 39 bilateral brain regions shared in both cohorts [44, 45]. To ensure a comparable age range across cohorts, we further restricted the HCP sample to participants aged 45-80 years. Therefore, the final analytic sample included 36,812 individuals (mean (SD) age = 63.41 (7.52) years; 54.95% female) from UKB and 391 individuals (mean (SD) age = 60.47 (9.64) years; 57.03% females) from HCP.

Because dMRI acquisition protocols and preprocessing pipelines differed between UKB and HCP, including the use of different multi-shell diffusion schemes and processing workflows (details described elsewhere [42, 46]), harmonization [47] was performed to derive comparable WM integrity measurements across cohorts while adjusting for age and sex.

We used the UKB cohort as the training dataset to develop the predictive model and applied the resulting model to the HCP cohort as an independent test dataset. Brain age was constructed using both MLR and UMLR approaches, implemented with conventional LASSO and unbiased LASSO, respectively. We evaluated systematic prediction bias for both methods using 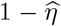, and assessed goodness of fit using *R*^2^ and RMSE.

We further calculated the brain age gap 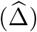, defined as predicted brain age minus chronological age. Negative 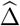 indicates younger and healthier appearing brain relative to chronological age [48], whereas larger positive 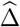 indicate accelerated brain aging, which may be related to neurodegeneration and cognitive decline.

To investigate the relationship between the brain-age gap (BAG) and cognitive performance, we assessed their association using the Fluid Cognition Composite Score in the HCP cohort [34]. Derived from the NIH Toolbox, this measure integrates performance across tests of reasoning, problem-solving and working memory, capturing cognitive functioning that is relatively independent of acquired knowledge. Higher scores indicate better cognitive performance. Accordingly, higher Fluid Cognition Composite Scores are generally associated with better brain health in older adults and may correspond to negative brain-age gaps. Therefore, the negative association estimated using UMLR-derived brain-age gaps may be more biologically plausible than the positive association obtained with conventional MLR-based estimates.

We further examined this relationship using *age-adjusted* fluid cognition. As shown in Table 1, both UMLR and MLR yielded negative associations, but only the association with the UMLR-derived BAG was statistically significant. The regression slope under UMLR was *−*0.170, which had a larger magnitude than the corresponding estimate of *−*0.074 under MLR. This result, together with the findings in Figure 4 of Case Study 1, is consistent with our theoretical result that 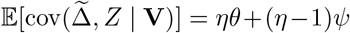. When age is not adjusted for, the term (*η −* 1)*ψ* can alter the direction of the association. When age is adjusted for, rendering (*η −* 1)*ψ* approximately zero, the association under MLR remains in the same direction as that under UMLR, but with a smaller magnitude (i.e., *−*0.074 under MLR versus *−*0.170 under UMLR).

**Table 1:** Association analysis of BAG with age-adjusted cognition composite score.

| Method | $\beta$ | S.E | $t$-value | $p$-value | $R^2$ | RMSE |
| --- | --- | --- | --- | --- | --- | --- |
| UMLR | -0.170 | 0.0504 | -3.38 | 0.0008 | 0.0310 | 15.8 |
| MLR | -0.074 | 0.0382 | -1.95 | 0.0520 | 0.0106 | 13.7 |

#### Case Study 2: Epigenetic clock

In this study, we used DNAm data measured in blood samples from the Framingham Heart Study Offspring cohort. DNAm levels at 443,206 CpG sites were assessed for 2,610 participants across two study sites. Specifically, 2,128 individuals were assayed at the University of Minnesota, and the remaining 482 Offspring samples were assayed at Johns Hopkins University.

Quality control (QC) and lab-specific normalization procedures were performed for both datasets [49, 50]. Briefly, QC included multidimensional scaling to identify sex mismatches and outliers, evaluation of sample and probe call rates, genotype concordance checks using overlapping single necleotide polymorphisms (SNPs), and removal of probes with ambiguous mapping or nearby SNPs. Samples and probes failing predefined criteria for missingness, concordance, or mapping were excluded (see [50] for further details).

We next applied sure independence screening [37] to reduce the number of CpG sites included in the MLR models. In addition, we retained CpG sites that were available in the FHS cohort and had been reported in previous epigenetic clock studies [11, 35, 36]. The final training dataset consisted of DNAm data for 2,990 CpG sites from 1,476 individuals in the UMN cohort (mean (SD) age = 65.12 (8.71) years; 54.40% female), and the testing dataset consisted of 226 individuals in the JHU cohort (mean (SD) age = 70.32 (8.50) years; 18.58% female). A harmonization procedure [47] was applied to DNAm data from the two laboratories, with adjustment for age and sex.

Epigenetic clocks were derived by both MLR and UMLR methods as in the brain age case study. In the downstream analysis, we examined the association between the epigenetic biological age gap and serum creatinine concentration (mg/dL), an index of kidney function that reflects creatinine clearance and glomerular filtration. Higher serum creatinine levels generally indicate reduced renal function and have been linked to adverse cardiometabolic outcomes and aging-related morbidity [51]. Accordingly, a biologically plausible expectation is a positive association between the epigenetic age gap and elevated serum creatinine. However, epigenetic clocks estimated using conventional MLR produced a negative association, likely driven by systematic prediction bias. In contrast, UMLR-derived epigenetic clocks yielded the expected positive association.

